# Inference of gene-environment interaction from heterogeneous case-parent trios

**DOI:** 10.1101/2022.10.15.512197

**Authors:** Pulindu Ratnasekera, Jinko Graham, Brad McNeney

**Author notes:** Correspondence: Brad McNeney.

## Abstract

In genetic epidemiology, log-linear models of population risk may be used to study the effect of genotypes and exposures on the relative risk of a disease. Such models may also include gene-environment interaction terms that allow the genotypes to modify the effect of the exposure, or equivalently, the exposure to modify the effect of genotypes on the relative risk. When a measured test locus is in linkage disequilibrium with an unmeasured causal locus, exposure-related genetic structure in the population can lead to spurious gene-environment interaction; that is, to apparent gene-environment interaction at the test locus in the absence of true gene-environment interaction at the causal locus. Exposure-related genetic structure occurs when the distributions of exposures and of haplotypes at the test and causal locus both differ across population strata. A case-parent trio design can protect inference of genetic main effects from confounding bias due to genetic structure in the population. Unfortunately, when the genetic structure is exposure-related, the protection against confounding bias for the genetic main effect does not extend to the gene-environment interaction term. We show that current methods to reduce the bias in estimated gene-environment interactions from case-parent trio data can only account for simple population structure involving two strata. To fill this gap, we propose to directly accommodate multiple population strata by adjusting for genetic principal components. We evaluate our approach through simulation and illustrate it on data from a study of genetic modifiers of cleft palate.

## 1 INTRODUCTION

We start by considering a log-linear model of population disease risk that includes main effects for genotypes *G*, environmental exposures *E*, and a gene-environment interaction term *G* × *E*. The *G* × *E* term allows genotypes to modify the effect of the exposure or, equivalently, the exposure to modify the effect of genotypes on the relative risk of developing the disease. Including a *G* × *E* term can improve model accuracy and provide a more detailed picture of disease etiology compared to models with just *G* and *E* main effects [9]. *G* × *E* is also useful for identifying environmental exposures with greater disease-association in individuals who carry particular alleles at susceptibility loci [23]. For example, dietary fat intake is more highly associated with obesity in carriers than in non-carriers of the Pro12Ala allele in the PPAR-γ gene [5].

We suppose throughout that *G* is an unmeasured causal locus in linkage disequilibrium with a measured non-causal test locus *G*′, and that the distribution of *GG*′ haplotypes differs across population strata (i.e. genetic structure). Stratum-specific differences in the *GG*′ haplotype frequencies can lead to differences in *G*′ risk across the population strata where none exist for G [29]. Exposure-related genetic structure occurs when the distribution of *E* also differs across the population strata [26]. Without some adjustment for the population strata, E will tag the stratum-specific differences in *G*′ risk (Figure 1), suggesting that *E* modifies *G*′ risk, even in the absence of *G* × *E* [18, 26]; we refer to this as spurious *G*′ × *E*.

**Figure 1.**
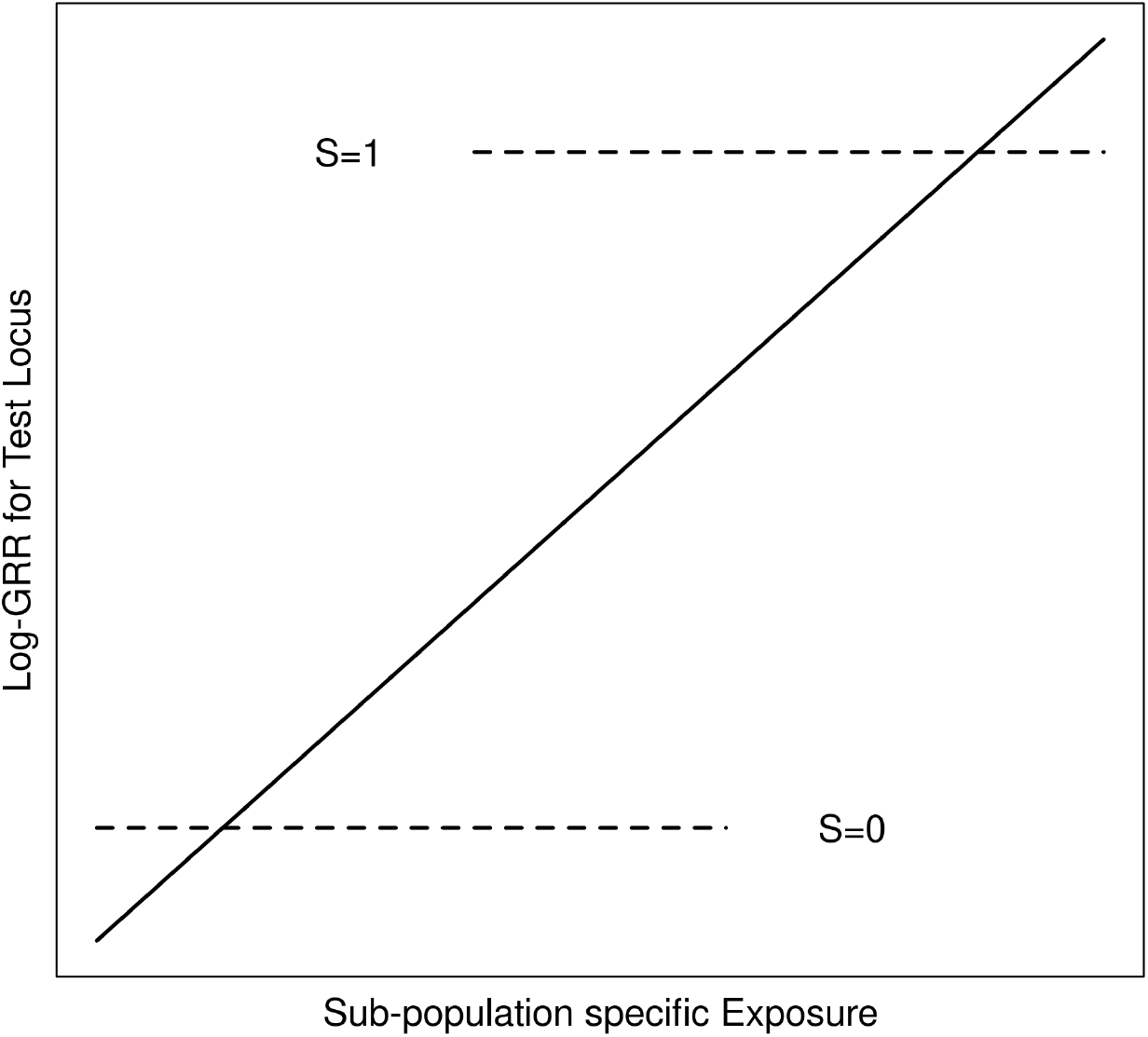
Schematic of log-GRRs for a non-causal test locus versus exposure in a structured population with two strata, S=0 and S=1. Dashed lines represent log-GRRs within each stratum. Horizontal positioning of these dashed lines indicates the support of the respective E distributions. High values of E are associated with S=1, in which one of the alleles at the test locus is associated with increased disease risk. Low values of E are associated with S=0 in which this same allele at the test locus is associated with low disease risk. Ignoring S yields the linear log-GRR curve indicated by the solid line, which erroneously suggests that E modifies the disease risk at the test locus.

A case-parent trio design can protect inference of genetic main effects from confounding bias due to genetic structure in the population [25]. In this design, investigators collect information on *G*′ and *E* in children affected with a disease of interest as well as the genotypes, 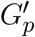, of their parents. To increase sample size, investigators may pool trios from multiple ancestral groups into one study; e.g., the GENEVA Oral Cleft Study [1] combined case-parent trios from recruitment sites in the United States, Europe and East Asia. Assuming *G*′ and *E* are independent within families, a log-linear model of disease risk leads to a conditional likelihood for the *G*′ and *G*′ × *E* effects, based on the child’s genotype given their exposure, affection status and parental genotypes [20]. Unfortunately, when the genetic structure is exposure related, the protection against confounding bias for the genetic main effect does not extend to the gene-environment interaction term [18, 26]. Thus, spurious *G*′ × *E* may be inferred from heterogeneous case-parent trio data in the absence of true *G* × *E*.

Methods to mitigate this bias may be classified as design-or data-based. For a binary environmental exposure, the *design*-based tetrad approach of [18] augments the case-parent trio by adding the exposure of an unaffected sibling. These authors control the bias by including the sibship-averaged exposure in the log-linear model. They show that all information about the interaction in the tetrad design comes from the siblings, not the parents [26]. Accordingly, they propose a sibling-augmented case-only design and analysis. By contrast, [20] takes a *data*-based approach, replacing the sibship-averaged exposure of [18] with the *predicted* exposure given ancestry. Predictions are obtained from a regression of exposure on principal components (PCs) computed from genetic markers that are unlinked to the test locus. This data-based approach may be applied to arbitrary exposures, including continuous exposures, and does not require siblings. However, its properties have not been evaluated in the case of more than two population strata.

We use the GENEVA Oral Cleft Study to motivate a new approach to unbiased inference of *G*′ × *E* in case-parent trios. The analysis of [2] found multiple single nucleotide polymorphisms (SNPs) that appeared to modify the effect of maternal smoking, maternal alcohol consumption or maternal multivitamin supplementation on the risk of cleft palate (CP). The self-reported ancestry of the study sample is primarily European or East Asian, and all three exposures are more common in self-reported Europeans than in self-reported East Asians [2, Table 2]. If the frequencies of haplotypes spanning causal SNPs also vary by ancestral groups, exposure-related genetic structure may lead to spurious gene-environment interaction. [13] focused on the self-reported Europeans and East Asians in the GENEVA Oral Cleft Study data. Applying the approach of [20], they confirmed the gene-environment interaction found by [2], and concluded that the evidence for gene-environment interaction is predominantly from the data of self-reported Europeans. These authors also considered whether exposure-related genetic structure *within* self-reported Europeans could explain the apparent *G*′ × *E*. Their results were inconclusive, however, possibly owing to the methodology’s limitation to just two ancestry groups. In modern datasets, the possibility of both inter- and intra-continental genetic structure necessitates methods that can more flexibly accommodate multiple ancestries. In this work we propose such an approach which relies on direct use of the genetic PCs to adjust for population structure.

**Table 1.** *GG*′ haplotype frequencies in four population strata.

| $GG'$ | Stratum | | | |
| --- | --- | --- | --- | --- |
| | $S = 0$ | $S = 1$ | $S = 2$ | $S = 3$ |
| R1 | 0.0 | 0.5 | 0.375 | 0.125 |
| R0 | 0.5 | 0.0 | 0.125 | 0.375 |
| N1 | 0.5 | 0.0 | 0.125 | 0.375 |
| N0 | 0.0 | 0.5 | 0.375 | 0.125 |

**Table 2.** Estimated type 1 error rates (top entry) and corresponding 95% confidence intervals (bottom entry) when data are simulated from 2, 3 or 4 strata with equal (top three rows) or unequal (bottom three rows) stratum sizes

| Equal stratum sizes |  |  |  |
| --- | --- | --- | --- |
| Adjustment | Number of strata |  |  |
|  | 2 | 3 | 4 |
| S | 0.0556<br>(0.049, 0.062) | 0.0524<br>(0.046, 0.0586) | 0.0498<br>(0.044, 0.056) |
| EEGM | 0.0538<br>(0.048, 0.060) | 1.0000<br>NA | 1.0000<br>NA |
| PC | 0.0546<br>(0.048, 0.061) | 0.0534<br>(0.047, 0.060) | 0.0496<br>(0.044, 0.056) |
| Unequal stratum sizes |  |  |  |
|  | 2 | 3 | 4 |
| S | 0.0524<br>(0.046, 0.058) | 0.0482<br>(0.042, 0.054) | 0.0536<br>(0.047, 0.059) |
| EEGM | 0.0536<br>(0.047, 0.060) | 1.0000<br>NA | 1.0000<br>NA |
| PC | 0.0540<br>(0.048, 0.060) | 0.0508<br>(0.045, 0.057) | 0.0527<br>(0.046, 0.059) |

The manuscript is structured as follows. In Section 2 we develop our direct PC-adjustment method and compare it to the indirect PC-based approach of [20]. In Section 3 we present simulations to evaluate the statistical properties of both approaches. In Section 4 we re-analyze the GENEVA data. Section 5 includes a discussion and areas for future work.

## 2 MODELS AND METHODS

### 2.1 Overview

We start with a log-linear model of disease risk parametrized in terms of genotype relative risks (GRRs) at a causal locus *G*. Under this model, *G* × *E* is equivalent to GRRs that depend on the exposure *E*. We then derive the GRRs at a non-causal test locus *G*′ in linkage disequilibrium with *G* and show that, in the absence of *G* × *E*, the *G*′-GRRs can depend on *E* when there is dependence between *E* and *GG*′ haplotypes in the population. Such dependence can lead to spurious inference of *G*′ × *E* in the absence of *G* × *E*. However, valid inference is obtained if we adjust the risk model for any variable *X* for which *E* and *GG*′ haplotypes are conditionally independent given *X* [20]. We review the rationale for the adjustment used by [20] in this context, and propose an alternative adjustment based on inferred population structure. In particular, we use the method of [6] to select a parsimonious set of PCs with which to adjust the risk model. A key question is whether the PC-selection method yields a set of PCs that provide enough adjustment to maintain type 1 error in the absence of *G* × *E*, but not so much that we compromise power in the presence of *G* × *E*. The Models and Methods section concludes with a discussion of the simulation methods used to answer this question.

### 2.2 Risk model and likelihood

Let *G* = 0, 1 or 2 denote the number of copies of the variant allele at the causal locus and E denote the exposure variable. The disease-risk model of [20] can be obtained from a log-linear model of the GRRs

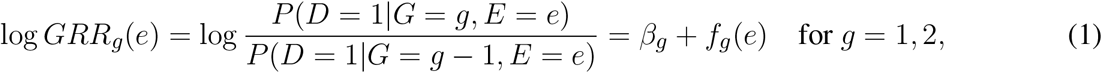

and the log-disease risk for carriers of the baseline genotype *G* = 0

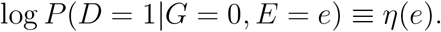

The parameters *β_g_* and *f_g_* (·) are, respectively, genotype-specific main effects and functions that allow for *G* × *E* interaction. We can also write disease risk in terms of the baseline risk *η*(*e*) and the GRRs as follows. First define *GRR*_0_(*e*) ≡ 1. Next, note that

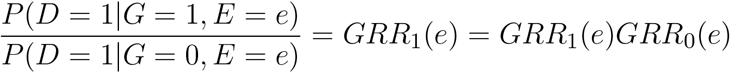

and

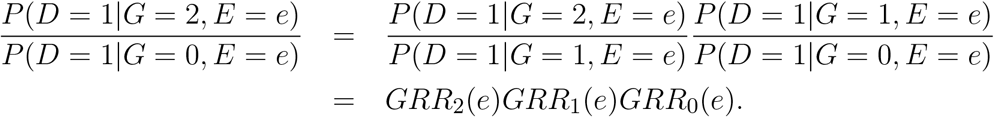

It follows that

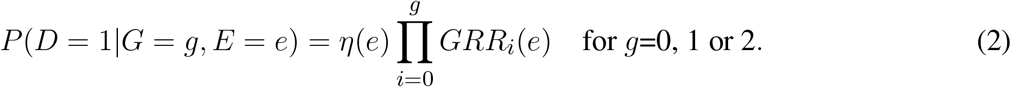

A likelihood for estimation of the GRR parameters *β_g_* and *f_g_*(·), *g* = 1, 2, from case-parent trio data can be derived under the assumption that *G* and *E* are conditionally independent given parental genotypes *G_p_*. As shown in Appendix A.1, the likelihood is based on the conditional probability of the child’s genotype given their exposure and parental genotypes. The function *η*(·) that parametrizes the environmental main effect drops out of the likelihood and cannot be estimated from case-parent trio data.

### 2.3 GRRs at a non-causal test locus

Let *G*′ denote genotypes at a non-causal test locus in linkage disequilibrium with the causal locus *G*. We assume *D* and *G*′ are conditionally independent given *G* and *E*, so that

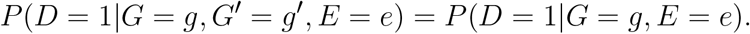

Therefore, the risk of disease given *G*′ and *E* can be written as

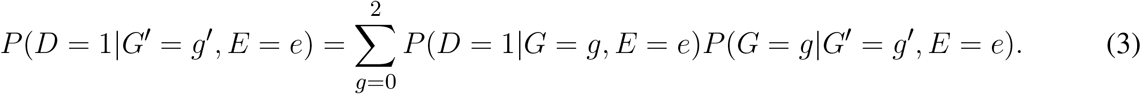

Equation (3) is a latent-class model [27] with the unobserved causal locus G as the latent class having probabilities *P*(*G* = *g*|*G*′ = *g*′, *E* = *e*). Equations (3) and (2) enable the log-GRRs at *G*′ to be written in terms of the latent-class probabilities and the GRRs at *G* as follows:

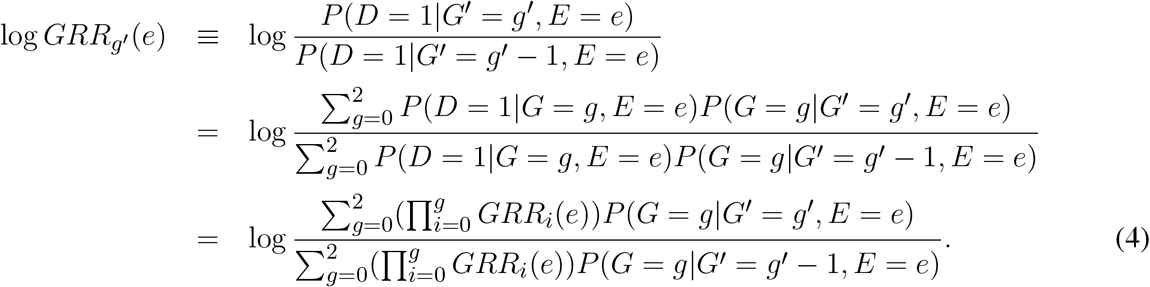

Without *G* × *E*, GRRs at *G* do not depend on *E*. Importantly, though, the log-GRRs at *G*′ *can* depend on *E* through the latent-class probabilities *P*(*G* = *g*|*G*′ = *g*′, *E* = *e*). In fact, as shown in Appendix A.2, these latent-class probabilities will depend on *E* whenever *GG*′ haplotypes and *E* are associated, as happens when the population has exposure-related genetic structure. Since *G*′ × *E* is equivalent to *GRR*_*g*′_ varying with *E*, equation (4) gives insight into how exposure-related genetic structure creates spurious *G*′ × *E*.

### 2.4 Augmented risk model

The development so far has considered a disease-risk model that depends only on *E* and a causal locus *G*. We now consider an augmented disease-risk model that depends on *E, G* and a third variable *X*:

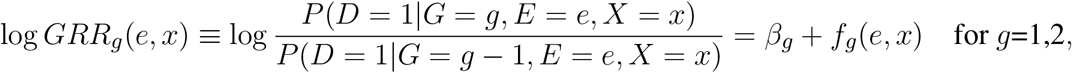

where *β_g_* and *f_g_*(·, *x*) are, respectively, genotype-specific main effects and functions that allow for *G* × *E* × *X* interaction. Defining

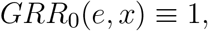

an analogous development to Section 2.3 leads to the following *X*-adjusted log-GRRs at *G*′:

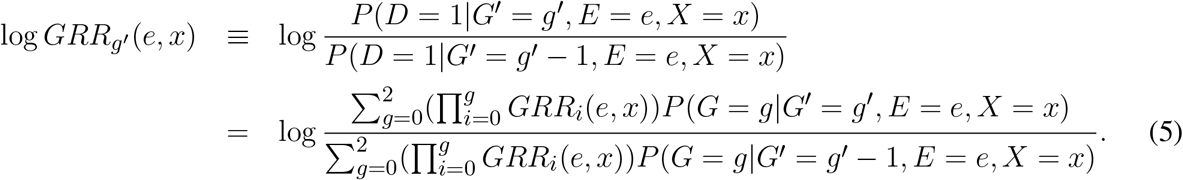

In the next section we discuss choices for *X* that eliminate *E* from the latent-class probabilities for *G*, and hence eliminate spurious *G*′ × *E* arising from exposure-related genetic structure.

### 2.5 Removing dependence of the latent-class probabilities on *E*

The diagram in Figure 2 depicts the dependence between *GG*′ haplotypes and *E* from exposure-related genetic structure in the population. In the figure, *S* is a categorical variable that indicates population strata. The categorical variable *X_E_* is a “coarsening” of *S* such that different levels of *X_E_* correspond to different *E* distributions, and, similarly, *X*_*GG*′_ is a coarsening of *S* such that different levels of *X*_*GG*′_ correspond to different *GG*′ haplotype distributions.

**Figure 2.**
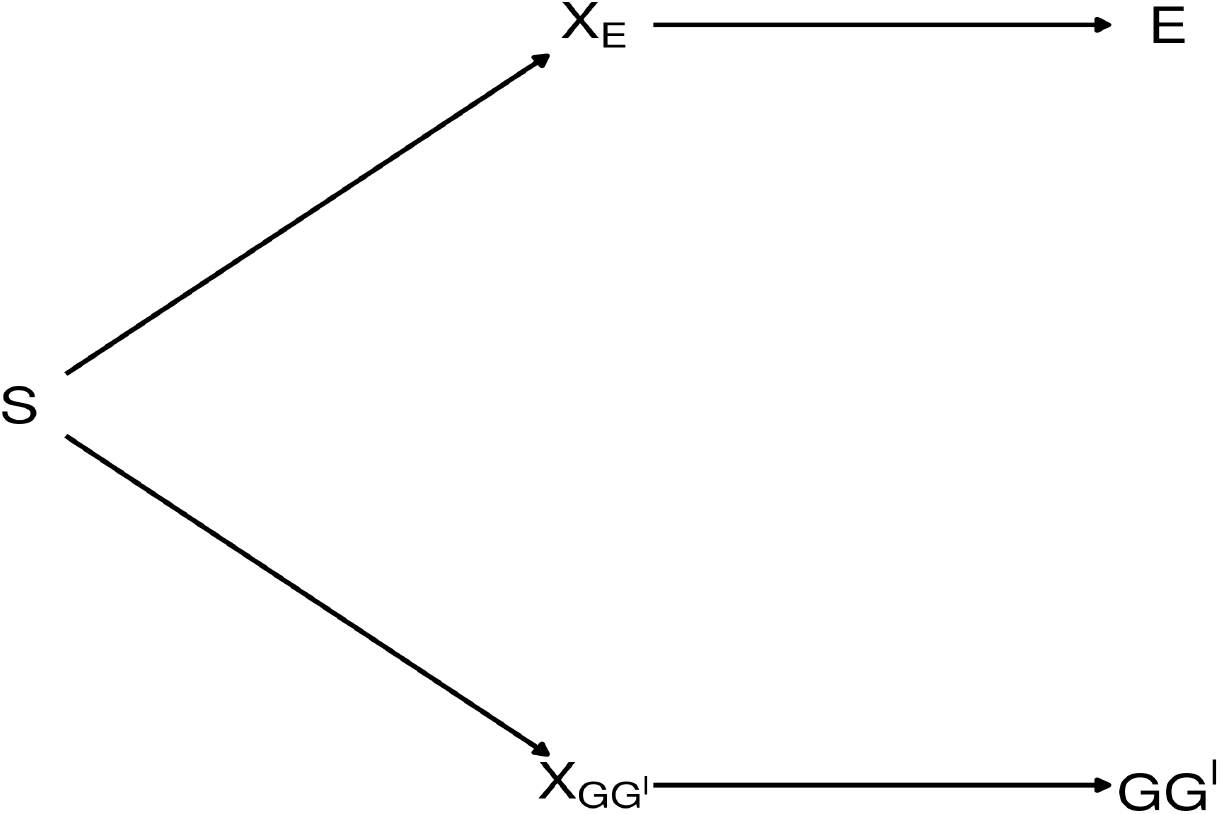
Diagram depicting exposure-related genetic structure. The latent population strata *S* induce dependence beetween *E* and *GG*′. Latent factors *X_E_* and *X*_*GG*′_ encode different distributions of *E* and *GG*′, respectively. *E* and *GG*′ are conditionally independent given any of the three variables that lie on the path between them.

The path connecting *E* and *GG*′ in Figure 2 is said to be *blocked* by each of the variables *X_E_, S* and *X*_*GG*′_ [12, Definition 1]. Therefore, *E* and *GG*′ are conditionally independent given any of the blocking variables *X_E_*, *S* or *X*_*GG*′_ [11]. As shown in Appendix A.2, a consequence is that conditioning on any of these variables removes the dependence of the latent-class probabilities on *E*. That is, letting *X* denote any of *X_E_*, *S* or *X*_*GG*′_, *P*(*G* = *g*|*G*′ = *g*′, *E* = *e*, *X* = *x*) = *P*(*G* = *g*|*G*′ = *g*′, *X* = *x*). Consequently, from equation (5),

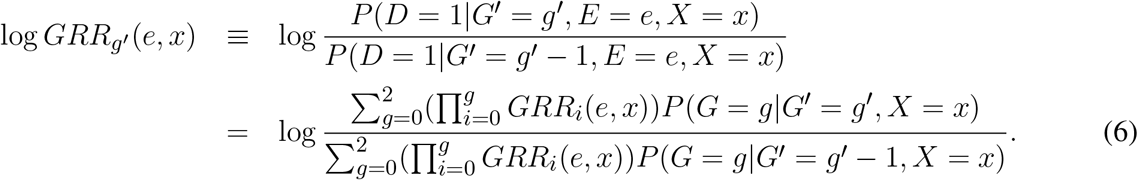

GRRs at *G*′ will thus depend on E if and only if GRRs at G do.

### 2.6 Linear model for the log GRRs

From equation (6) we see that, for fixed *g*′ and *x*, log *GRR*_*g*′_(*e, x*) varies with *e* if and only if the *GRR_g_*(*e, x*) do. We can therefore test for *G* × *E* by fitting a model for log*GRR*_*g*′_(*e, x*) that allows separate curves in *e* for each combination of *g*′ and *x* [21]. We take these curves to be straight lines, and test whether any of them have non-zero slope. For a fixed value *x* of the adjustment variable *X* and a fixed value *e* of the environmental exposure *E*, the log-GRR is:

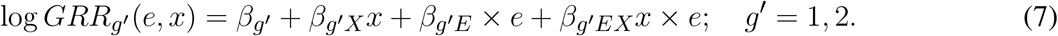

The generalization of the above model to a vector *X* is to replace *β*_*g*′*X*_*x* with 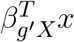 and *β*_*g*′*EX*_ with 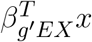 for coefficient vectors *β*_**g*′*_ and *β*_*g*′*EX*_. The intercepts of the log-GRR curves, *β*_*g*_′ + *β*_*g*′*X*_*x*, are the genetic main effects in stratum *x* (i.e. when *e* = 0). The slopes, *β*_*g*′*E*_ + *β*_*g*′*EX*_*x*, are the *G*′ × *E* interaction terms in stratum *x*. We use a likelihood-ratio test of the null hypothesis that *β*_*g*′*E*_ = *β*_*g*′*EX*_ = 0 for *g*′ = 1, 2, *versus* the alternative hypothesis that at least one of these slope parameters is non-zero to detect *G* × *E*. We emphasize that the simplified log-GRR curves in e characterize *G* × *E* rather than environmental main effects, which are not estimable from case-parent trio data. Genetic main effects *are* estimable however and flexibly parametrized by the intercept terms of the log-GRR curves. The flexibility in the intercept terms avoids mis-specification of the genetic main effects which can lead to biased inference of interaction effects [28].

### 2.7 Choice of *X*

Following [18], [20] set *X* to be the categorical variable *X*E** that distinguishes E distributions among the genetic strata of the population. Since *X_E_* is unobserved, [20] consider the expectation of *E* given genetic markers (EEGM) as a surrogate 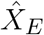. The idea behind their EEGM approach is to distinguish exposure distributions by their mean, which may vary across genetic strata, *S*. Though *S* is not known, it is reflected in principal components (PCs), *Ŝ*, computed from a set of genetic markers that are unlinked to *G*′. The expectation of *E* given *Ŝ* can be estimated by linear regression of *E* on *Ŝ* when *E* is continuous, or by logistic regression when *E* is binary. For EEGM adjustment, the expected exposure within genetic strata is estimated by 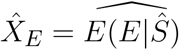. [20] showed that EEGM adjustment works well where there are two population strata, but our simulation results (Section 3) indicate that it works poorly for more than two strata. We therefore propose to adjust for population strata directly; i.e., to take *X* = *S*. In particular, if the population has *K* + 1 genetic strata, indexed 0,…, *K*, we let *S* denote a vector of *K* dummy variables that distinguish these strata such that that the *k*th element *S_k_* = 1 for trios in stratum *k* > 0 and 0 otherwise, for *k* = 1,…, *K*.

### 2.8 Inferred population strata

The population stratum variable *S* reflects genetic ancestry and is not generally known. Since adjustment for self-reported ancestry can lead to bias [24] we use marker-based PCs, *Ŝ*. An advantage of PC-adjustment is that it does not enforce discrete strata, and individuals whose PC values lie between those of clusters on the PC plot (e.g. admixed individuals) will have intermediate values of the slope and intercept of their log-GRR curve.

Standard PC adjustment in genetic association analyses relies on a relatively large set of PCs. For *K* PCs the degrees of freedom of the test for *G*′ × *E* is equal to 2(*K* + 1). Thus, using more PCs than are necessary reduces the power of the test for *G*′ × *E*. We seek methods to select a parsimonious set of PCs that provides enough adjustment to control type 1 error rate, without sacrificing power. We consider three PC-selection methods. The first [30] is an automated version of the graphical approach of looking for an “elbow” in the scree plot of variance explained by the PCs as a function of their number. The second [6] is an estimator of the rank of a matrix under a model in which the data matrix is a noisy version of a low-rank matrix. The third [10] is to select PCs corresponding to eigenvalues that exceed a significance threshold determined from the distribution of the largest eigenvalue of an unstructured random matrix.

### 2.9 Simulation methods

#### 2.9.1 Simulating *G*, *G*′ and *E* on case-parent trios

To study the statistical properties of our proposed approach and compare it to the method of [20], we generated 5000 data sets of 3000 informative case-parent trios. Trios were sampled from one of four population strata labelled *S* = 0, 1, 2 or 3. We assumed random mating within and no mixing between strata. We performed some simulations using equal-sized strata and others using unequal-sized strata. In the case of unequal stratum sizes, the split was 60%, 40% for two strata; 50%, 30% and 20% for three strata; and 40%, 30%, 20% and 10% for four strata.

For a given stratum, informative trios were simulated following the methods proposed by [22, 21]. Briefly, *GG*′ haplotypes are first simulated on parents in a random-mating population according to the stratum-specific *GG*′ haplotype distributions in Table 1. Child haplotypes are then simulated following Mendel’s laws and assuming no recombination between *G* and *G*′. The child’s exposure *E* is also simulated according to the stratum-specific distributions described below. Finally, the child’s disease status is simulated based on the disease-risk model (1). Trios with an affected child and at least one heterozygous parent at the test locus are retained. The data recorded on each trio are 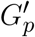, *G*’, and *E*, where 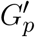 is the pair of parental genotypes at the test locus.

Spurious *G*′ × *E* is induced by specifying different distributions of E and *GG*′ haplotypes in the four strata of Table 1. The *GG*′ distributions for strata *S* = 0 and *S* = 1 are as in [20]. Alleles at *G* are denoted *R* (risk) and *N* (non-risk), while alleles at *G*′ are denoted 1 and 0. We summarize the haplotype distributions by the implied allelic correlations between the index alleles *R* and 1. Under the *GG*′ haplotype frequencies given in Table 1, these correlations are *r*_0_ = −1 in stratum *S* = 0, *r*_1_ = 1 in stratum *S* = 1, *r*_2_ = 0.5 in stratum *S* = 2 and *r*_3_ = −0.5 in stratum *S* = 3.

The stratum-specific distributions of *E* are chosen to be normal with common variance *σ*^2^ = 0.36, and means *μ*_0_ = −0.8, *μ*_1_ = 0.8, *μ*_2_ = 2.4 and *μ*_3_ = 4.0 in strata 0, 1, 2 and 3, respectively. The *E* distributions for strata *S* = 0 and *S* = 1 are as in [20].

The disease-risk model is specified as follows. The genetic main effect is *β_g_* = log(3)/2 for *g* = 1, 2, corresponding to a 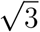-fold increase in relative risk for each copy of the risk allele (R) in the absence of *G* × *E*.To evaluate the type 1 error rate of the *G* × *E* test we set *f_g_*(*e*) = 0 in our simulations. To investigate power we choose a linear interaction model for the *G* × *E* term, setting *f_g_*(*e*) = *β_gE_e* with *β_gE_* = −0.10, −0.15, −0.20 or −0.25.

#### 2.9.2 Simulating markers for PC adjustment

A standard method of PC adjustment is to calculate PCs from a genomic region that is unlinked to the test locus. It is recommended that markers in this region be thinned, or LD pruned, to have pairwise correlations of *r*^2^ ≤ 0.1 [7]. We simulated such panels of markers based on data from the 1000 genomes project [4] using two East Asian (Chinese Dai in Xishuangbanna, China [CDX] and Han Chinese in Bejing China [CHB]) and two European (Iberian population in Spain [IBS] and Finnish in Finland [FIN]) populations. From the initial download of the genome-wide data, we retained 6,929,035 diallelic, autosomal markers with minor allele frequency (MAF) 0.05 or greater in all four of the population groups.

Our initial approach to simulating markers for a given population stratum was to fit a hidden Markov model (HMM) to the haplotypes in that stratum, chromosome by chromosome, using fastPHASE [16], and use this fitted model to simulate individual multilocus genotypes using SNPknock [17]. The simulated data are then LD pruned and principal components are computed from the thinned panel of markers. However, the computation involved in this approach proved to be prohibitive. For example, fitting the HMMs took up to 5 hours per chromosome. We therefore considered two computationally cheaper alternatives. In the first alternative, we started from an LD-pruned set of markers in the original data and fit HMMs to this set. In the second alternative, we used the same panel of pruned markers, but simulated genotypes *independently* based on the MAFs in the population strata. In what follows we refer to the first and second alternatives as *LD-based* and *independent* marker simulation, respectively.

Independent markers could contain more information about the population strata than markers in LD. As a result, PC adjustment with independent markers might control type 1 error more effectively than adjustment with markers in LD. To assess this possibility, we completed 100 preliminary simulation replicates using LD-based marker simulation and 5000 replicates using independent marker simulation. We simulated trios from four population strata under the null hypothesis of no *G* × *E*, used the PC selection method of [6] to adjust the risk model and estimated the resulting type 1 error rates. Estimated type 1 error rates and their 95% confidence intervals under the LD-based and independent simulation methods were 0.04 (0.002, 0.078) and 0.0496 (0.044, 0.056), respectively, and consistent with similar type 1 error rates for the two approaches. We therefore used the faster simulation of independent markers for the simulation study.

In Sections 3.2 and 3.3 we present type I error and power results for two, three or four population strata. For two strata (*S* = 0 and *S* = 1), marker simulations were based on the CHB and IBS population groups. For three strata (*S* = 0, *S* = 1 and *S* = 2), simulations were based on the CHB, IBS and CDX population groups.

## 3 RESULTS

### 3.1 Selection of Principal Components

All PC selection methods performed well when the sizes of the population strata were equal (results not shown), but not when the sizes were unequal. We illustrate with simulation results involving datasets of 3000 trios sampled from four unequal-sized strata. For *K* + 1 = 4 populations we require *K* = 3 PCs. In 5000 simulation replicates, the method of [6] always selected three, the method of [30] always selected one, and the method of [10] selected three PCs 4942 times and four PCs 58 times. Other simulation results with unequal-sized strata (not shown) yielded similar results. Therefore, in what follows we use the method of [6] to select PCs.

### 3.2 Type I Error Rate

We compared the type I error rates of the test for *G*′ × *E* using (i) adjustment with the true stratum membership *S*, (ii) the EEGM adjustment of [20], and (iii) PC adjustment. Results for simulated datasets with equal or unequal stratum sizes are shown in Table 2. For both equal and unequal stratum sizes, adjustment by *S* or direct PCs maintains the nominal 5% error rate regardless of the number of strata. By contrast, EEGM adjustment leads to an inflated type I error rate when there are more than two strata. In light of the inflated size of the test, we do not consider EEGM adjustment in the following section on power.

### 3.3 Power

Table 3 provides a comparison of estimated power when data are simulated from two, three or four strata. Results are shown for simulations using both equal and unequal stratum sizes and for different values of the *G* × *E* effect. From these results we see that power increases with effect size, decreases with number of strata and tends to be slightly larger for unequal strata than equal strata. Importantly, the estimated power under PC adjustment is always within simulation error of that under adjustment for true stratum membership.

**Table 3.** Estimated power (top entry) and corresponding 95% confidence intervals (bottom entry) of different adjustment schemes for different *G* × *E* interaction effects *β_gE_*, number of strata and stratum-size distributions.

|  |  | Equal stratum sizes |  |  |  |
| --- | --- | --- | --- | --- | --- |
| | | $\beta_{gE}$ | | | |
| Num. Strata | Adjustment | -0.10 | -0.15 | -0.20 | -0.25 |
| 2 | S | 0.2602<br>(0.248, 0.272) | 0.5660<br>(0.552, 0.580) | 0.8420<br>(0.832, 0.852) | 0.9558<br>(0.950, 0.961) |
|  | PC | 0.2580<br>(0.246, 0.270) | 0.5660<br>(0.552, 0.580) | 0.8404<br>(0.830, 0.850) | 0.9564<br>(0.951, 0.962) |
| 3 | S | 0.1742<br>(0.164, 0.185) | 0.3844<br>(0.371, 0.398) | 0.6498<br>(0.636, 0.663) | 0.8288<br>(0.818, 0.839) |
|  | PC | 0.1788<br>(0.168, 0.189) | 0.3920<br>(0.378, 0.406) | 0.6616<br>(0.648, 0.675) | 0.8316<br>(0.821, 0.842) |
| 4 | S | 0.1306<br>(0.121, 0.140) | 0.2766<br>(0.264, 0.289) | 0.5010<br>(0.487, 0.515) | 0.6970<br>(0.684, 0.710) |
|  | PC | 0.1396<br>(0.130, 0.149) | 0.2936<br>(0.281, 0.306) | 0.5088<br>(0.495, 0.523) | 0.6918<br>(0.679, 0.704) |
|  |  | Unequal stratum sizes |  |  |  |
| | | $\beta_{gE}$ | | | |
|  |  | -0.10 | -0.15 | -0.20 | -0.25 |
| 2 | S | 0.2636<br>(0.251, 0.276) | 0.5724<br>(0.559, 0.586) | 0.8328<br>(0.822, 0.843) | 0.9518<br>(0.946, 0.958) |
|  | PC | 0.2648<br>(0.252, 0.277) | 0.5722<br>(0.558, 0.586) | 0.8322<br>(0.822, 0.842) | 0.9514<br>(0.945, 0.957) |
| 3 | S | 0.1950<br>(0.184, 0.206) | 0.4322<br>(0.418, 0.446) | 0.7082<br>(0.696, 0.721) | 0.8640<br>(0.854, 0.874) |
|  | PC | 0.1936<br>(0.183, 0.204) | 0.4334<br>(0.420, 0.447) | 0.7054<br>(0.693, 0.718) | 0.8632<br>(0.854, 0.873) |
| 4 | S | 0.1614<br>(0.151, 0.172) | 0.3470<br>(0.334, 0.360) | 0.6028<br>(0.589, 0.616) | 0.7894<br>(0.778, 0.801) |
|  | PC | 0.1598<br>(0.150, 0.170) | 0.3380<br>(0.325, 0.351) | 0.5872<br>(0.574, 0.601) | 0.7820<br>(0.770, 0.794) |

## 4 THE GENEVA ORAL CLEFT STUDY

### 4.1 Data and objectives

The GENEVA Oral Cleft study [1] is comprised of 550 case-parent trios from 13 different sites across the United States, Europe, Southeast and East Asia. Data were obtained through dbGAP at https://www.ncbi.nlm.nih.gov/projects/gap/cgi-bin/study.cgi?study_id=phs000094.v1.p1 with accession number phs000094.v1.p1. Of the 550 trios, only 462 were available for analysis. Summaries of the trios by ancestry and gender of the affected child are shown in Table 4. From this table we see the ancestry of the sample is predominantly European (46%) and East Asian (51%).

**Table 4.**
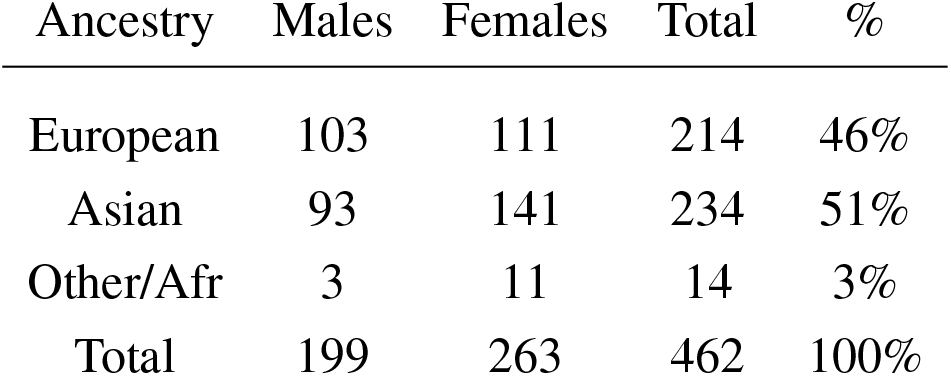
Gender of 462 affected children by self-reported ancestry

| Ancestry | Males | Females | Total | % |
| --- | --- | --- | --- | --- |
| European | 103 | 111 | 214 | 46% |
| Asian | 93 | 141 | 234 | 51% |
| Other/Afr | 3 | 11 | 14 | 3% |
| Total | 199 | 263 | 462 | 100% |

The objective of the GENEVA study is to discover genetic contributions to orofacial clefts, the most common type of craniofacial birth defect in humans, and to assess whether these genes modify the effect of exposures known to be associated with cleft palate. Maternal exposure to multivitamins, alcohol and smoking were assessed through maternal interviews focused on the peri-conceptual period (3 months prior to conception through the first trimester), which includes the first 8-9 weeks of gestation when palatal development is completed. Exposure status is summarized in Table 5. From this table we see that the three dichotomous exposures are all more common in Europeans. In contrast to the continuous exposures of the simulation study, the exposures we consider in the GENEVA study are all dichotomous.

**Table 5.** Exposure rates for maternal alcohol consumption, maternal smoking and maternal vitamin supplementation by self-reported ancestry in affected trios.

| Ancestry | Percent exposed to Maternal |  |  | Affected children |
| --- | --- | --- | --- | --- |
|  | Alcohol | Smoking | Vitamin Supp. |  |
| European | 41% | 28% | 57% | 214 |
| East Asian | 4% | 3% | 21% | 234 |
| Other/Afr | 14% | 7% | 71% | 14 |
| Total | 21% | 14% | 39% | 462 |

### 4.2 GENEVA data analysis

#### 4.2.1 PC selection

LD pruning of the genome-wide panel of SNPs at an *r*^2^ threshold of 0.1 yielded 63,694 markers. In a principal component analysis of these markers, the first PC explains 6.3% of the total variance and all others explain less than 0.4%. Not surprisingly, the method of [6] selects one PC. A plot of the projections of the data onto the first two PCs is shown in Figure 3, with points colored by self-reported ancestry. Each PC has been shifted by subtracting the minimum value and scaled by the range so that the values are between zero and one. The first PC distinguishes those with self-reported East Asian ancestry from those with self-reported European ancestry; hence, a value near zero corresponds to a hypothetical East Asian and a value near one corresponds to a hypothetical European. The second PC separates the single self-reported African child from all others.

**Figure 3.**
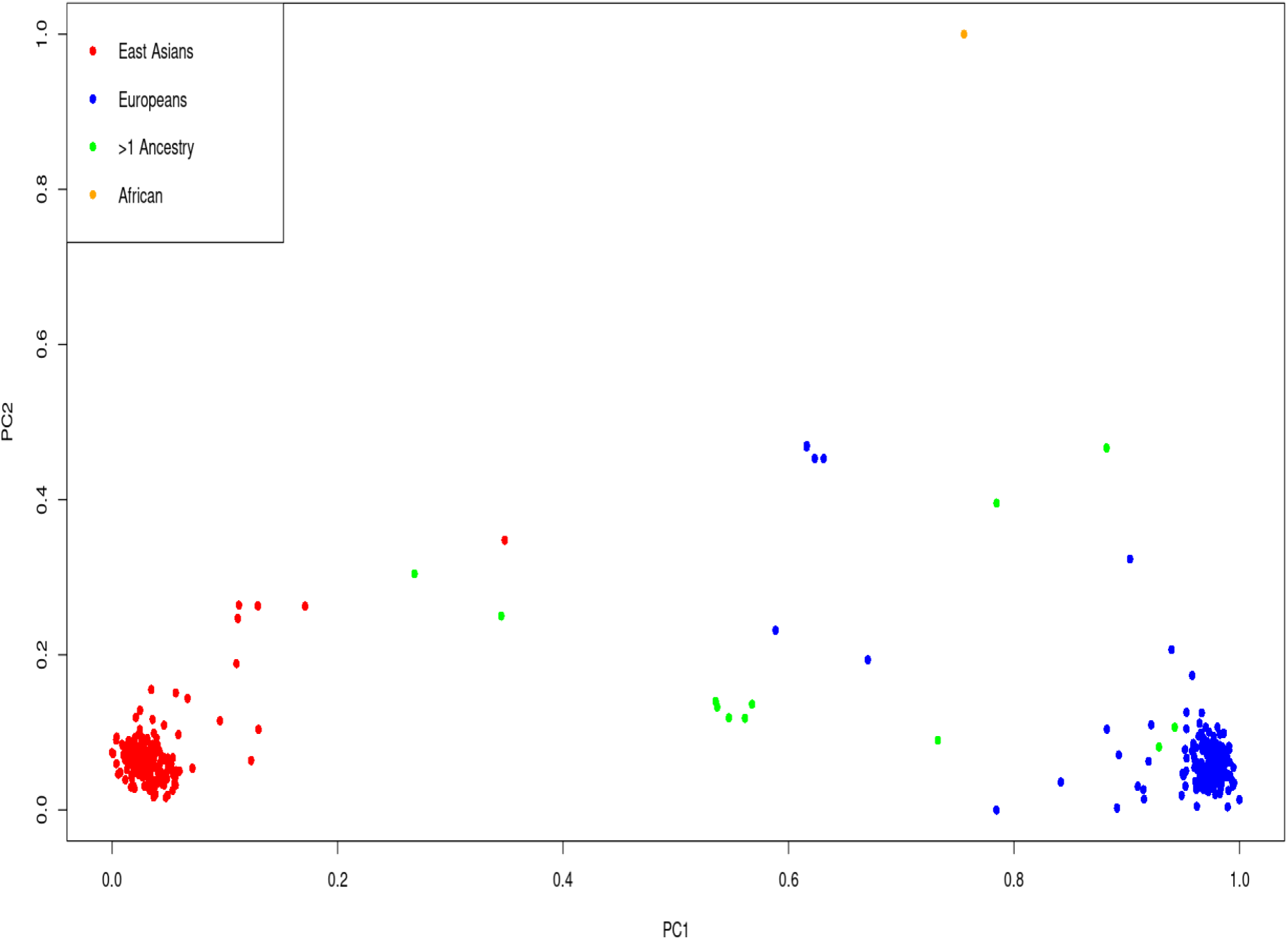
Projections of each affected child onto the first two PCs by self-reported ancestry: red=East Asian (234 trios), blue=European (214 trios), orange=African (one trio) and green=multiple ancestry/other (13 trios). Each PC has been shifted and scaled so that a PC1 value near zero corresponds to a hypothetical East Asian and a PC1 value near one corresponds to a hypothetical European.

#### 4.2.2 Inference of *G* × *E*

The conditional-likelihood methods outlined in Appendix A.1 were applied to the data. We focused on inference of *G* × *E* between maternal alcohol consumption and the six SNPs in the *MLLT3* gene that had significant *G* × *E* at the 5% level in the analysis of [2]. Displays of the LD between these SNPs and others nearby [19] are shown in Figure S1, Appendix A.3, for self-reported European subjects and self-reported East Asian subjects. Table 6 shows the results of fitting three different log-linear models of *G*′ × *E*. Following [2], each is based on an additive genetic model that specifies equal log-GRRs for genotypes *g*′ = 1 or 2. Results based on fitting a more general co-dominant model (1) were similar (results not shown). The first model, as in [2], makes no adjustment for exposure-related genetic structure in the population, the second uses EEGM adjustment and the third uses PC adjustment. From the table we see that, for each test SNP, p-values for the tests of *G*′ × *E* are smallest when we make no adjustment. Comparing the EEGM and PC adjustment approaches we find that p-values from PC adjustment are similar to, but tend to be slightly smaller than, those from the EEGM adjustment. Of the six test SNPs show in the table, four retain significance at the 5% level after adjustment for exposure-related genetic structure.

**Table 6.**
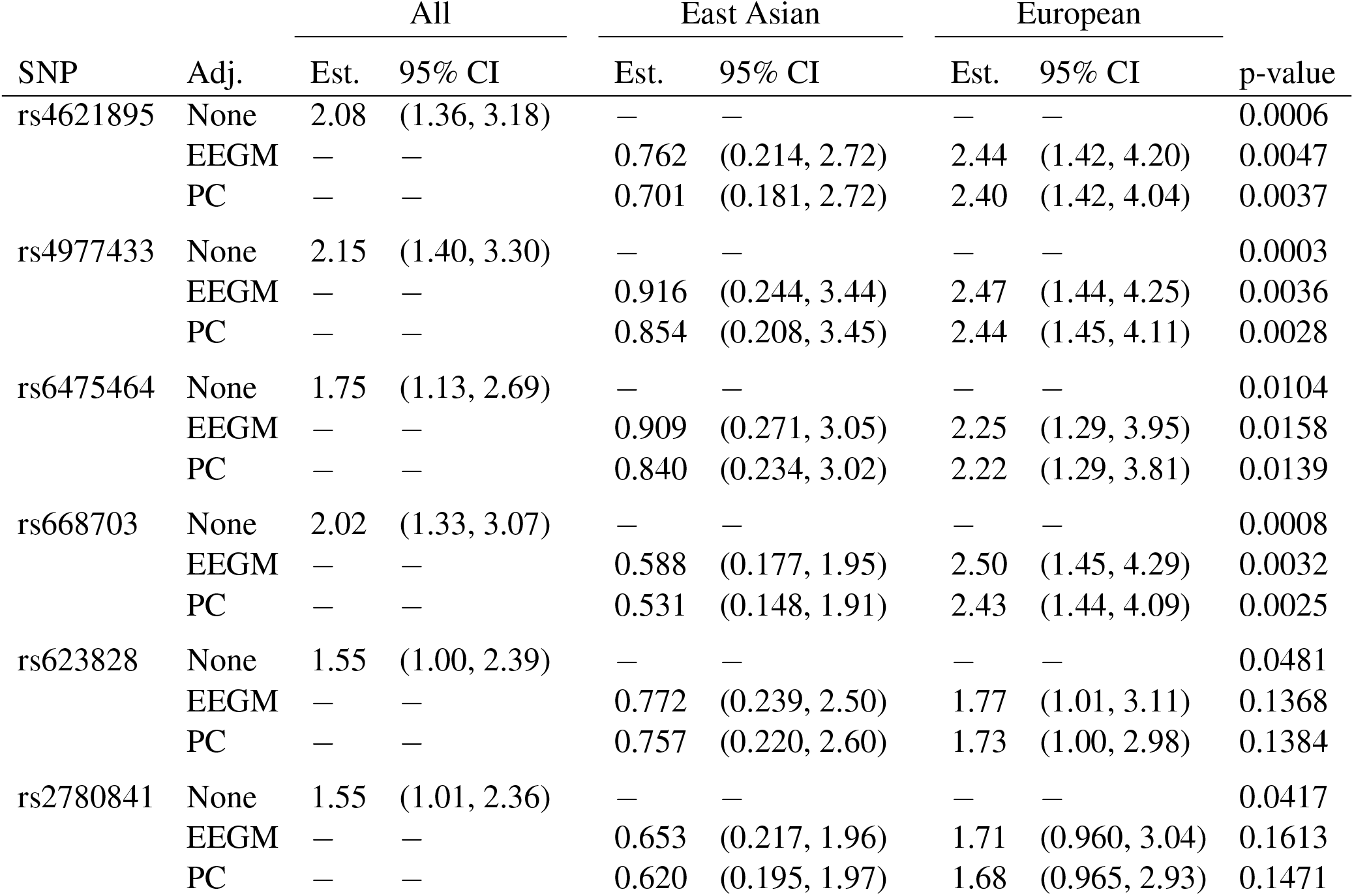
Estimated modifying effects of maternal alcohol consumption on GRRs, 95% confidence intervals and p-values from the analysis of the GENEVA data, at 6 SNPs in the MLLT3 gene (Chr 9) showing significant interaction with maternal alcohol consumption in [2]. Estimates, confidence intervals and tests are based on fitting an additive genetic model and use (i) no adjustment, (ii) EEGM adjustment or (iii) PC adjustment to control for exposure-related genetic structure in the population. The unadjusted analysis considers all trios without regard to genetic structure. The EEGM- and PC-adjusted analyses allow for genetic structure and we have reported estimates for hypothetical East Asian and European subjects.

The estimates shown in Table 6 are of the multiplicative factors by which maternal alcohol consumption modifies the GRRs at the six test SNPs. For a binary exposure such as maternal alcohol consumption, these modifying effects can be obtained by exponentiating the interaction term in the log-GRR model. With no adjustment for genetic structure there is a single interaction term and hence a single estimated modifying effect for all trios. For example, maternal alcohol consumption is estimated to increase the GRR at SNP rs4621895 by a factor of about 2.1 for all trios. By contrast, with EEGM or PC adjustment the interaction term depends on the value of the adjustment variable and we have reported estimates for hypothetical East Asian and European subjects in our sample. For example, maternal alcohol consumption is estimated to decrease the GRR at SNP rs4621895 by a factor of about 0.73 for East Asian trios and to increase the same GRR by a factor of about 2.4 for European trios. For these data, the adjustment variables used in the EEGM- and PC-adjustment approaches are highly correlated (Pearson correlation 0.996), and so the estimates for the two approaches are very similar. These estimates are also similar to those obtained from an analysis using self-reported ancestry (results not shown). The 95% confidence intervals for hypothetical East Asians cover one for each SNP but do not cover one for hypothetical Europeans, with the exception of SNP rs2780841. These results suggest that any *G* × *E* signal is from trios of European ancestry, where maternal alcohol consumption is more common.

## 5 DISCUSSION

We consider a log-linear model of GRRs at a causal locus G. Under this model, *G* × *E* is equivalent to GRRs that vary with the exposure E. We show that exposure-related genetic structure in the population can lead to spurious *G*′ × *E* at a non-causal test locus *G*′ in LD with *G*. However, valid inference of *G*′ × *E* can be obtained by augmenting the GRR model with a blocking variable *X*, such that *GG*′ haplotypes and *E* are conditionally independent given *X*. We discuss the choice of *X* for inference of *G*′ × *E* when data are collected from a study of case-parent trios. The population strata *S* would be an ideal choice for *X* but may not be known definitively. We propose to use principal components (PCs) instead. In particular, we calculate PCs from a genomic region unlinked to the test locus and select a parsimonious subset using the method of [6]. We then specify a linear model for the log-GRRs whose intercept and slope depend on PC values. Slopes that vary with PC values allow the modifying effect of the exposure to vary with population strata, which can be important for maintaining power [20, Section 3.3]. Through simulations, we show that our PC adjustment maintains the nominal type-1 error rate and has nearly identical power to detect *G* × *E* as an oracle approach based directly on *S*. We illustrate our approach by applying it to an analysis of real data from case-parent trios in the GENEVA Oral Cleft Study. In our analysis of the GENEVA data, we focussed on SNPs and exposures identified by [2]. In a discussion of their results, these authors noted that the SNPs they identified are not in known cleft-palate susceptibility genes and are either intronic or are upstream/downstream of coding regions. This lack of compelling biological plausibility, coupled with the striking differences in exposure distributions between the self-reported European and East Asian strata, motivated our *G* × *E* analysis that adjusts for population structure. However, our results (Table 6) and those of [13] do not contradict the hypothesis of *G* × *E*, but rather suggest that any *G* × *E* signal is due to the self-reported European trios. Further data collection aimed at self-reported European trios may provide stronger conclusions regarding the presence of *G* × *E*.

To reduce bias from exposure-related genetic structure, direct PC adjustment has advantages over the EEGM approach and design-based strategies such as the tetrad approach of [18] and the sibling-augmented case-only approach of [26]. Unlike the EEGM approach, PC adjustment controls the type 1 error when there are more than two population strata. Unlike the design-based strategies, PC adjustment does not require siblings nor assume binary exposures.

Development of alternative approaches based on propensity scores is an area for future work. The EEGM approach is attractive in that it reduces the genetic principal components to a single score, *E*(*E*|*Ŝ*). For binary exposures, such as those in the GENEVA study, the EEGM is a propensity score [15]. For continuous exposures, such as those in the simulation study, the analog to the EEGM is a continuous-treatment propensity score [3]. With continuous exposures, we could predict *E* given the genetic markers and *then* convert the predictions to a Normal density score that takes low values for predictions far from their observed value. These density scores could be used either as predictors [8] or weights [14] in subsequent analyses. It would be interesting to explore the use of propensity-score methods in inference of *G*′ × *E* from case-parent trios with continuous exposures, particularly when there are more than two population strata.

## AUTHOR CONTRIBUTIONS

PR developed the statistical methods, performed the simulations and data analyses, and wrote the initial draft of the manuscript. BM and JG conceptualized the study and revised the manuscript. All authors proofread and approved the final version of the manuscript.

## ACKNOWLEDGMENTS

Funding support for the study entitled “International Consortium to Identify Genes and Interactions Controlling Oral Clefts” was provided by several previous grants from the National Institute of Dental and Craniofacial Research (NIDCR). Data and samples were drawn from several studies awarded to members of this consortium. Funding to support original data collection, previous genotyping and analysis came from several sources to individual investigators. Funding for individual investigators include: R21-DE-013707 and R01-DE-014581 (Beaty); R37-DE-08559 and P50-DE-016215 (Murray, Marazita) and the Iowa Comprehensive Program to Investigate Craniofacial and Dental Anomalies (Murray); R01-DE-09886 (Marazita), R01-DE-012472 (Marazita), R01-DE-014677 (Marazita), R01-DE-016148 (Marazita), R21-DE-016930 (Marazita); R01-DE-013939 (Scott). Parts of this research were supported in part by the Intramural Research Program of the NIH, National Institute of Environmental Health Sciences (Wilcox, Lie). Additional recruitment was supported by the Smile Train Foundation for recruitment in China (Jabs, Beaty, Shi) and a grant from the Korean government (Jee).

The genome-wide association study, also known as the Cleft Consortium, is part of the Gene Environment Association Studies (GENEVA) program of the trans-NIH Genes, Environment and Health Initiative [GEI] supported by U01-DE-018993. Genotyping services were provided by the Center for Inherited Disease Research (CIDR). CIDR is fully funded through a federal contract from the National Institutes of Health (NIH) to The Johns Hopkins University, contract number HHSN268200782096C. Funds for genotyping were provided by the NIDCR through CIDR’s NIH contract. Assistance with genotype cleaning, as well as with general study coordination, was provided by the GENEVA Coordinating Center (U01-HG-004446) and by the National Center for Biotechnology Information (NCBI).

We sincerely thank all of the patients and families at each recruitment site for participating in this study, and we gratefully acknowledge the invaluable assistance of clinical, field and laboratory staff who contributed to this effort over the years.

## DATA AVAILABILITY STATEMENT

Datasets used for the analyses described in this manuscript were obtained from dbGaP at https://www.ncbi.nlm.nih.gov/projects/gap/cgi-bin/study.cgi?study_id=phs000094.v1.p1 through dbGaP accession number phs000094.v1.p1.

---

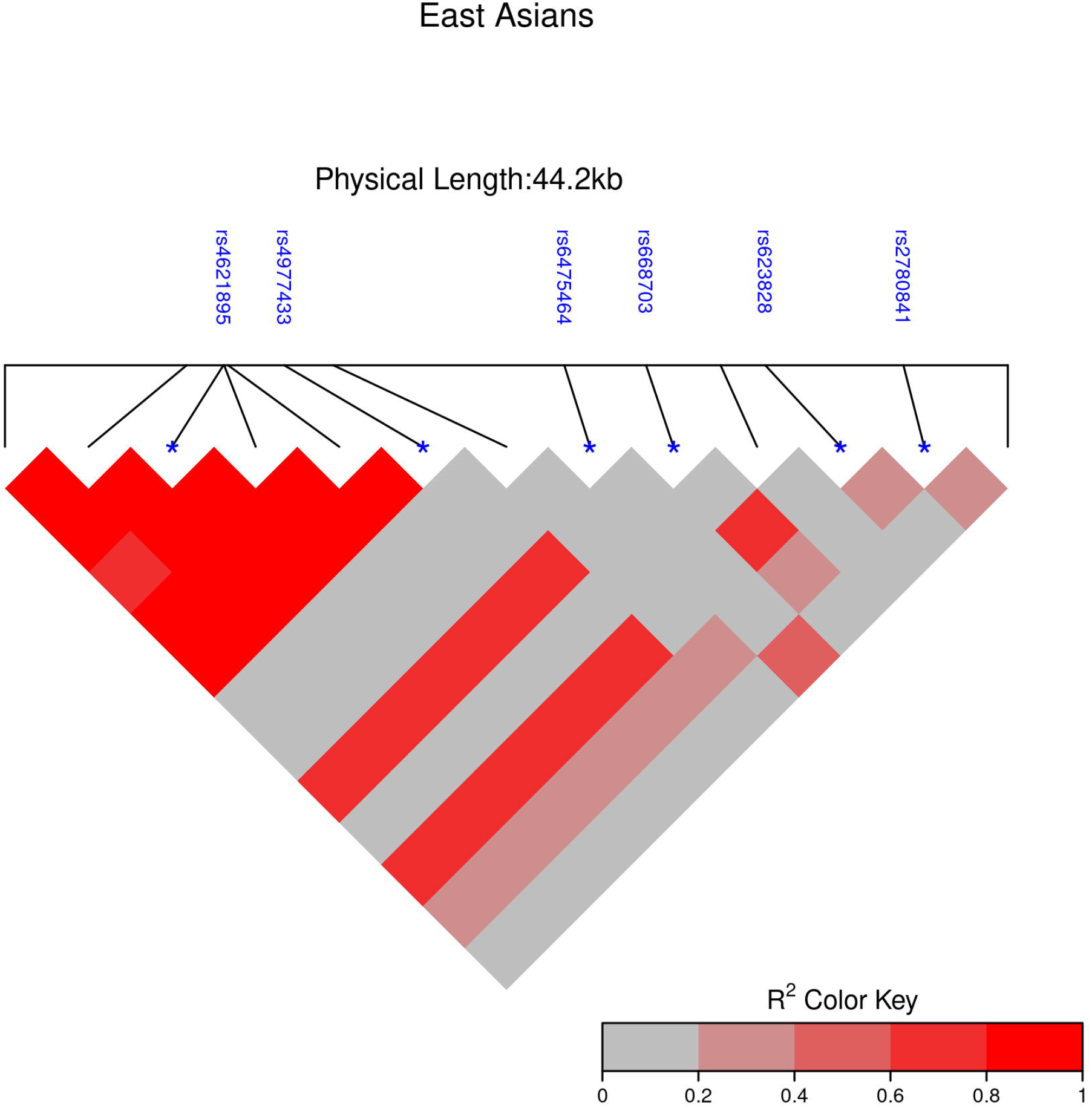

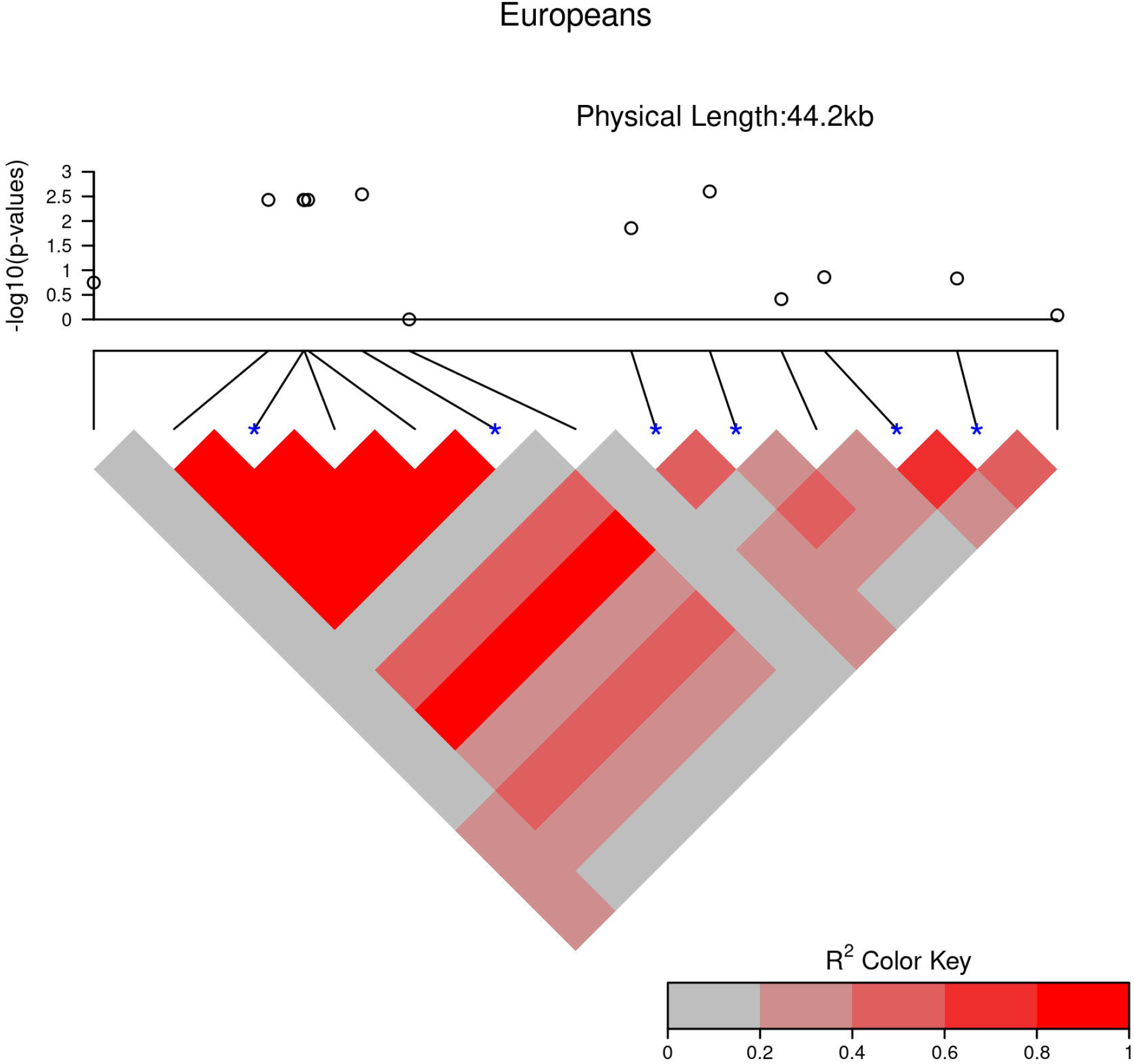

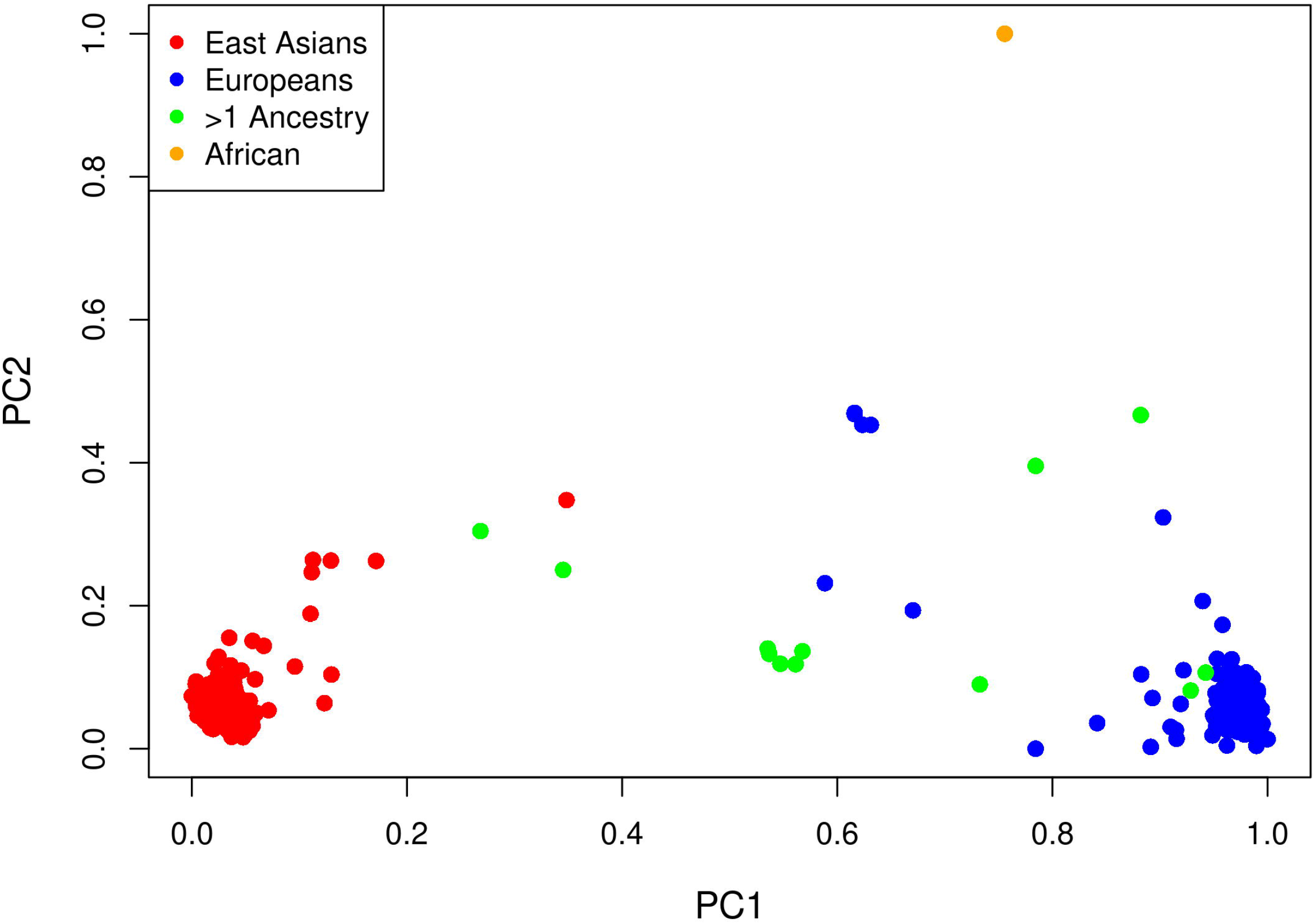

