## appendix for "Inference of gene-environment interaction from heterogeneous case-parent trios"

### Supplementary Material

#### A APPENDIX

##### A.1 Conditional likelihood and analysis

Assuming $G$ and $E$ are independent within families, one can write the conditional probability of the affected child's genotype given $E$ and $G_p$ in terms of the GRRs of equation (1), $GRR_g(e) = \exp(\beta_g + f_g(e))$. For example, when both parents are heterozygous, denoted below as $G_p = 3$ Shin et al. (2014), one can show that the child's genotype probabilities are:

$$P(G = 0 \mid D = 1, E = e, G_p = 3) = \frac{1}{1 + 2\exp(\beta_1 + f_1(e)) + \exp(\beta_1 + f_1(e) + \beta_2 + f_2(e))},$$

$$P(G = 1 \mid D = 1, E = e, G_p = 3) = \frac{2\exp(\beta_1 + f_1(e))}{1 + 2\exp(\beta_1 + f_1(e)) + \exp(\beta_1 + f_1(e) + \beta_2 + f_2(e))}, \quad \text{and}$$

$$P(G = 2 \mid D = 1, E = e, G_p = 3) = \frac{\exp(\beta_1 + f_1(e) + \beta_2 + f_2(e))}{1 + 2\exp(\beta_1 + f_1(e)) + \exp(\beta_1 + f_1(e) + \beta_2 + f_2(e))}.$$

A complete list of conditional genotype probabilities for the affected child is given in Table 1 of Shin et al. (2014). For an additive model, in which $\beta_1 = \beta_2 \equiv \beta$ and $f_1(e) = f_2(e) \equiv f(e)$, the model simplifies considerably; e.g.,

$$P(G = g \mid D = 1, E = e, G_p = 3) = \frac{\exp(O_g + g(\beta + f(e)))}{\sum_{i=0}^{2} \exp(O_i + i(\beta + f(e))},$$

where $O_g$ is an "offset" term that equals $\log 2$ for $g = 1$ and 0 otherwise.

The likelihood is a product of conditional probabilities over all trios in the study, viewed as a function of the parameters $\beta_1$, $\beta_2$, $f_1(e)$ and $f_2(e)$. Each trio's contribution to the likelihood can be viewed as the contribution of a matched set to a likelihood for a conditional logistic regression, in which the matched set comprises the affected child and other possible offspring of the parents, referred to here as the affected child's pseudo-siblings. After constructing appropriate matched sets, software for conditional logistic regression may be used to maximize the likelihood from a case-parent trio study.

Code in the R environment for statistical computing R Core Team (2022) is available to perform such analyses and may be obtained from the first author upon request. The code sets up a data frame with rows for each affected child and pseudo-sibling, and columns specifying the ID for each trio (`ID`), affection status coded as 1 for the affected child and 0 for pseudo-siblings, an offset variable (`O`) coded as $\log 2$ for a heterozygous offspring of doubly-heterozygous parents and 0 otherwise, and the $G$, $E$ and PC variables. We then call `clogit()` from the `survival` package Therneau (2021) to perform the conditional logistic regression. The argument to `clogit()` is a formula that specifies affection status as the response, trio IDs as `strata(ID)`, offsets as `offset(O)` and the other model terms. For an additive model, the other model terms are a main effect for $G$, two-way interactions between $G$ and $E$ and between $G$ and the PCs, and, finally, a three-way interaction between $G$, $E$ and the PCs.

#### A.2 Dependence of latent-class probabilities on $E$

Write the probabilities in terms of the conditional distribution of $GG'$ given $E$ as

$$P(G = g|G' = g', E = e) = \frac{P(G = g, G' = g'|E = e)}{\sum_{i=0}^{2} P(G = i, G' = g'|E = e)}.$$

Supposing that the numerator and denominator both depend on $E$, so may their ratio. However, if we condition on the blocking variable $X$

$$\begin{aligned} P(G = g|G' = g', E = e, X = x) &= \frac{P(G = g, G' = g'|E = e, X = x)}{\sum_{i=0}^{2} P(G = i, G' = g'|E = e, X = x)} \\ &= \frac{P(G = g, G' = g'|X = x)}{\sum_{i=0}^{2} P(G = i, G' = g'|X = x)} \\ &= \frac{P(G = g, G' = g'|X = x)}{P(G' = g'|X = x)} \\ &= P(G = g|G' = g', X = x). \end{aligned}$$

Thus, latent-class probabilities in the model adjusted for $X$ do not depend on $E$.

#### A.3 LDheatmaps of SNPs in *MLLT3*

LDheatmaps of pairwise $R^2$ values in and around the six SNPs in the *MLLT3* gene that showed significant $G \times E$ with maternal alcohol consumption in Beaty et al. (2011) are shown in Figure S1 for self-reported Europeans and self-reported East Asians. There is generally stronger pairwise LD between SNPs that showed significant $G \times E$ in the self-reported Europeans than in the self-reported East Asians. The $-\log_{10}$ p-values from the PC-adjusted analysis are shown above the self-reported Europeans, who appear to be the drivers of the $G \times E$ signal.

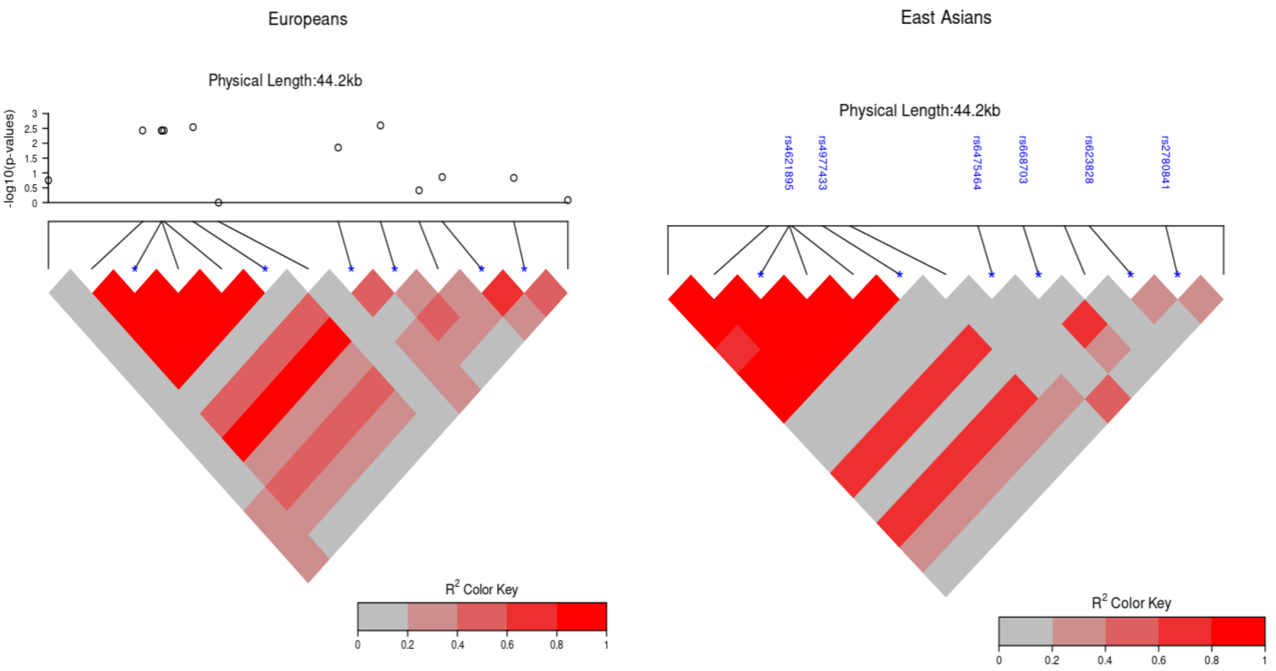


**Figure S1.** LDheatmap of pairwise $R^2$ values in and around the six SNPs in the *MLLT3* gene that showed significant $G \times E$ with maternal alcohol consumption in Beaty et al. (2011). Left panel: self-reported Europeans, with p-values from the PC-adjusted analysis shown above. Right panel: self-reported East Asians, with the names of the six SNPs shown above.
